# Investigating 3D microbial community dynamics in the rhizosphere using complex-field and fluorescence microscopy

**DOI:** 10.1101/2024.02.13.578483

**Authors:** Oumeng Zhang, Reinaldo E. Alcalde, Haowen Zhou, Siyuan Yin, Dianne K. Newman, Changhuei Yang

## Abstract

Microbial interactions in the rhizosphere contribute to soil health. Understanding these interactions thus has the potential to advance sustainable agriculture, ecosystem management, and environmental stewardship. Yet it is difficult to understand what we cannot see; amongst the limitations in rhizosphere imaging are challenges associated with rapidly and non-invasively imaging microbial cells over field depths relevant to plant roots. Here, we present a novel bimodal imaging technique called Complex-field and Fluorescence microscopy using the Aperture Scanning Technique (CFAST) that begins to address these limitations by integrating quantitative phase and 3D fluorescence imaging. We showcase CFAST’s practicality and versatility in two ways. First, by harnessing its depth of field of more than 100 microns, we significantly reduce the number of captures required for 3D imaging of plant roots and bacteria in the rhizoplane, thereby minimizing potential photobleaching and phototoxicity. Second, by leveraging CFAST’s phase sensitivity and fluorescence specificity, we track early bacterial aggregate development, bacterial competition, and gene expression under varying environmental conditions. Specifically, we resolve bacterial growth dynamics of mixed populations at the early stages of colonization without the need for genetically labeling environmental isolates. Moreover, we find that the expression of genes of interest to rhizosphere chemistry (e.g. representative genes involved in phosphorus-sensing and antibiotic production) varies spatiotemporally within microbial populations that are surface-attached and appears distinct from their expression in planktonic cultures. Together, CFAST’s attributes overcome commercial imaging platform limitations and enable new insights to be gained into microbial behavioral dynamics in experimental systems of relevance to the rhizosphere.

## Introduction

The nuanced biological processes within the rhizosphere (the zone of the soil in the vicinity of plant roots) play a crucial role in nutrient cycling, plant health, soil carbon dynamics, water relations, and microbial diversity (1–7). Advanced imaging methods, including confocal microscopy and molecular imaging techniques, enable real-time visualization, spatial mapping, and quantitative analysis of microbial activities around plant roots in a lab environment. They have the potential to track changes over time, aiding in the identification of microbial populations, biofilm structures, and nutrient gradients (8–13). Additionally, they can provide molecular and functional insights, such as real-time bacterial gene expression within the rhizosphere microbiome, but such applications are few and far between (14) due to complex geometry and dynamic interactions between roots and bacteria that are difficult to image even in a lab environment.

For example, fluorescence imaging offers exceptional specificity for differentiating components within the rhizosphere, yet to capture their full three-dimensional (3D) dynamics, traditional fluorescence microscopy often necessitates axial scanning. This approach slows down the imaging process and increases the risks of photobleaching and phototoxicity. Recently, the ability to resolve the 3D fluorescence distribution without scanning the sample has been made possible through advances in point spread function (PSF) engineering (15–17) combined with computational methods (18–20). These methods involve the modulation of light at the pupil of a microscope, thereby altering the detected image to encode 3D information within a 2D image. Another challenge in fluorescence imaging is that labeling is not always a viable option for environmental samples. Hence, a non-invasive imaging approach would be particularly valuable for imaging the rhizosphere. Quantitative phase imaging emerges as a less explored yet promising alternative in this context. This technique captures the complex optical field as light passes through the sample. Despite its lower specificity compared to fluorescence imaging, it offers distinct advantages such as its high sensitivity when visualizing low contrast samples. In contrast to most phase imaging techniques, such as quantitative differential phase contrast (21, 22), digital holography (23–25), and Fourier ptychographic microscopy (26–28), the synthetic aperture imaging based on Kramers-Kronig relations (KKSAI) (29) adopts simple pupil manipulation. Therefore, it is compatible with PSF engineering to offer complementary information when imaging microbial communities associated with plant roots.

In this study, we present Complex-field and Fluorescence microscopy using the Aperture Scanning Technique (CFAST), a novel bimodal imaging approach designed to overcome the aforementioned limitations to better study plant-microbe and microbe-microbe interactions in a lab setting. CFAST represents a bimodal approach merging 3D fluorescence and complex field imaging, delivering uniform resolution over an extended depth of field. Here we demonstrate the capability of CFAST to provide detailed, non-invasive images of rhizosphere organisms across different length scales. While these experiments represent a first step towards imaging the rhizosphere *in situ*, more generally, CFAST holds promise as an instrument through which to gain insight into biological dynamics in various complex environments.

## Results

### System schematic and operating principle

The schematic of CFAST is illustrated in Figure 1a. A 4f system (L1 and L2) is integrated into a conventional wide-field microscope to modulate light at the pupil of the objective lens. The system uses a plane wave from a laser to illuminate the sample and excite the fluorophores simultaneously. A spinning disk (SD) aperture, the heart of this design, is positioned at the back focal plane (BFP) of the 4f system, blocking three-quarters of the pupil. The spinning disk stops at four specific orientations [Figure 1a (i-iv)] during the acquisition of fluorescence and brightfield images. A dichroic mirror (DM) subsequently separates the fluorescence and illumination light, which are then captured by two cameras.

**Figure 1.**
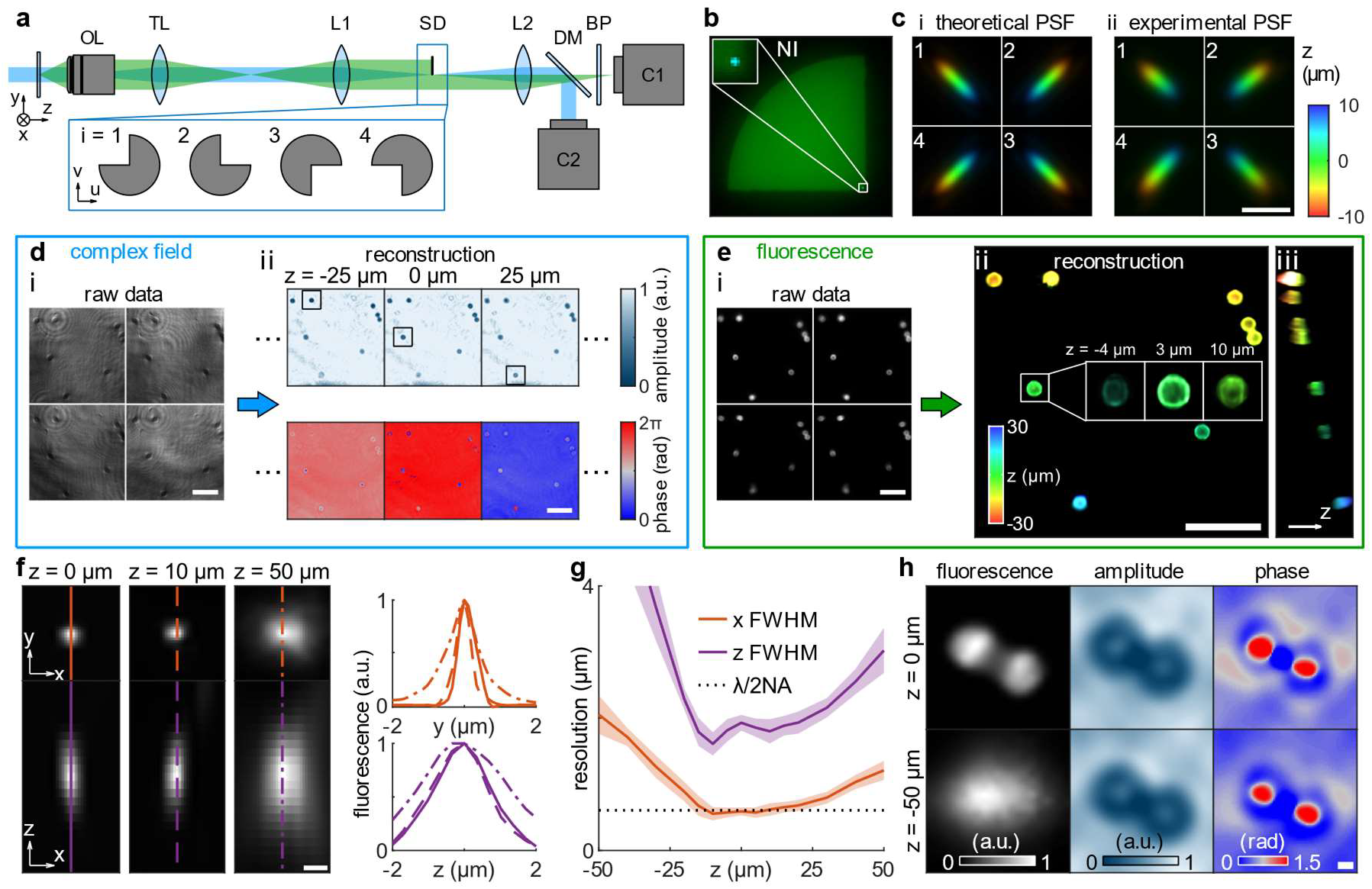
Schematic and working principle of Complex-field and Fluorescence microscopy using Aperture Scanning Technique (CFAST). (a) OL, objective lens; TL, tube lens; L1 and L2, 4f lenses; SD, spinning disk; DM, dichroic mirror; BP, bandpass filter. Four brightfield images and four fluorescence images are captured by detectors C1 and C2, respectively, at different orientations of SD (*i* ∈ {1,2,3,4}). Blue represents the illumination laser; green represents the fluorescence. (b) Image of the back focal plane when SD is oriented as depicted in (a) (*i* = 1). The illumination laser is aligned such that the normal incident (NI) component is at the corner of the SD. (c) (i) Theoretical and (ii) experimental point spread functions within a depth of field of 20 microns. (d) Workflow of phase imaging. (i) Four raw images of 5-micron (radius) fluorescent beads on a tilted cover glass are analyzed using the complex field reconstruction algorithm, generating a z stack of the amplitude and phase images. (e) Workflow of fluorescence imaging. (i) Four raw images of the aforementioned sample are analyzed using a 3D deconvolution algorithm (see Methods), generating (ii,iii) fluorescence images of the whole 3D volume. (ii) The xy view; (iii) the yz view. (f) The xy and xz cross section and the corresponding line profiles of the 3D fluorescence reconstruction of a 100-nm bead at different axial positions. (g) The (orange) lateral and (purple) axial resolution of the fluorescence imaging in CFAST is determined using the full width at half maximum (FWHM) of the line profiles. Lines and shaded areas represent the average and standard deviation across 63 beads. (h) Fluorescence, amplitude, and phase reconstruction of two 1.25-micron beads that are (top) in focus and (bottom) defocused by 50 microns. Scale bar, 5 microns in (c); 50 microns in (d,e); 1 micron in (f,h).

Synthetic aperture imaging based on Kramers-Kronig relations (KKSAI) (29) is utilized for complex field imaging. The aperture is aligned such that the normal incidence (NI) component of the illumination laser passes through its corner (Figure 1b). This unscattered NI component and the portion of the scattered light fields that transmit through the aperture will interfere with each other at the image plane, and the resulting image will be detected by one of the cameras. Briefly, the Kramers-Kronig (KK) relations establish bidirectional connections between the real and imaginary components of a complex function that is analytic in the upper half plane (30, 31). In KKSAI, the (shifted) complex optical field *S*_*i*_ of the opening of the spinning disk, termed sub-aperture, corresponding to each detected image *I*_*i*_ (*i* ∈ {1,2,3,4}) can be written as (29)

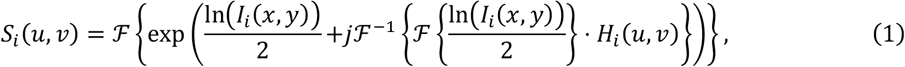

where (*u, v*) and (*x, y*) are the coordinates at the pupil and image planes, *ℱ* is the Fourier transform, and *H*_*i*_(*u, v*) represents a Hilbert kernel that depends on the sub-aperture position (see Methods for more details). After shifting and stitching the fields of four sub-apertures, the phase and amplitude of the sample are obtained from the complete optical field of the pupil through a Fourier transform.

We experimentally demonstrated KKSAI by imaging 5-micron radius fluorescent beads on a tilted cover glass (Figure 1d). Four brightfield images [*I*_*i*_, Figure 1d(i)] are captured to compute *S*_*i*_ according to Equation (1). Although the recovered complex optical field [amplitude and phase, Figure 1d(ii)] is 2D, it enables us to digitally generate a z-stack by introducing a defocusing phase to the pupil (32, 33). For example, the sharpest focus of the top, middle, and bottom beads within the boxed sections in Figure 1d(ii) corresponds to axial positions of -26, -3, and 22 microns, respectively. This indicates an approximate cover glass tilt of 15°.

The principle of three-dimensional fluorescence imaging resembles the multi-view reflector microscope (34) and the Fourier light-field microscope (35). Given that only part of the pupil gets imaged in each of the four frames, the images corresponding to a point source exhibit apparent lateral shifts as the source defocuses (Supplementary Figure S1); each frame has a unique shift direction. The theoretical and experimental PSFs are shown in Figure 1c. The 3D fluorescence imaging procedure is shown in Figure 1e; four fluorescence images of the aforementioned fluorescent beads [Figure 1e(i)] are analyzed using a modified Lucy-Richardson deconvolution algorithm(35–37) (see Methods). The reconstructed 3D images [Figure 1e(ii,iii)] confirm the cover glass tilt, consistent with the KKSAI results (Figure 1d). The deconvolution algorithm is also validated using synthetic data (Supplementary Figure S2), where CFAST outperforms other widely-used 3D PSFs (15, 38).

To quantify the resolution of 3D fluorescence imaging in CFAST, we imaged 100-nm fluorescent beads at various axial positions and measured the full width at half maximum (FWHM) of the xy and xz cross-sections of the reconstruction (Figure 1f). The experimental resolution along the lateral and axial directions are 0.6 ± 0.1 and 1.9 ± 0.1 microns for in-focus beads, respectively (Figure 1g). The lateral resolution is consistent with the theoretical diffraction limit and remains consistent within a 20-micron depth of field. The axial resolution shows a slight enhancement with a defocus of ±10 microns. Notably, this depth of field significantly outperforms the specification of the objective lens (depth of field = 1.6 microns, 20×, 0.42 NA). Further, compared to the 20-micron depth of field of 3D fluorescence imaging, KKSAI shows a more remarkable depth of field. For instance, when two 1.25-micron fluorescent beads are ∼3 microns apart, the 3D fluorescence algorithm fails to recover both spheres when defocused by 50 microns, whereas complex field images still resolve the separation (Figure 1h).

The bimodal imaging provided by CFAST offers numerous advantages. In the following sections, we show applications with plant roots and bacteria to highlight that (1) the long depth of field of complex field imaging (more than 100 microns) improves the efficiency of 3D fluorescence imaging, and (2) quantitative phase imaging provides an additional dimension that complements fluorescence imaging (e.g., Supplementary Figure S3).

### Bimodal imaging of plant roots and bacteria on the rhizoplane with varying depth of field

Standard scanning techniques, such as confocal imaging, are typically slow when capturing 3D volumes. Moreover, given the intricate nature of plant roots, their thickness often varies considerably across different sections. Therefore, scanning a target region without a prior understanding of its thickness can result in suboptimal scan speeds and risk overexposing the specimen to excitation light, potentially accelerating photobleaching and phototoxicity. CFAST offers a solution to this limitation by leveraging the long depth of field of KKSAI; the thickness of the sample can be determined by obtaining amplitude and phase images at just one axial position.

We demonstrate this ability of CFAST by imaging the 3D morphology of maize roots through its autofluorescence (Figure 2a,b and Supplementary Figure S4). In a thin section of the maize root, the recovered amplitude [Figure 2a(i)] shows both edges with sharp focus simultaneously, eliminating the need of additional axial positions to capture the full 3D fluorescence distribution [Figure 2a(ii-v)]. In contrast, for a thick region of the same root, KKSAI indicates a 70-micron axial difference between its left and right sections [Figure 2b(i)]. Considering the depth of field of 20 microns in 3D fluorescence imaging, we deduce that capturing at least four axial positions is essential for the full 3D volume reconstruction [Figure 2b(ii-v)]. Fourier ring correlation (FRC) (39) analysis show lateral resolutions of 0.9 to 1.2 microns in the thin sections and 1.2 to 1.3 microns in the thick sections.

**Figure 2.**
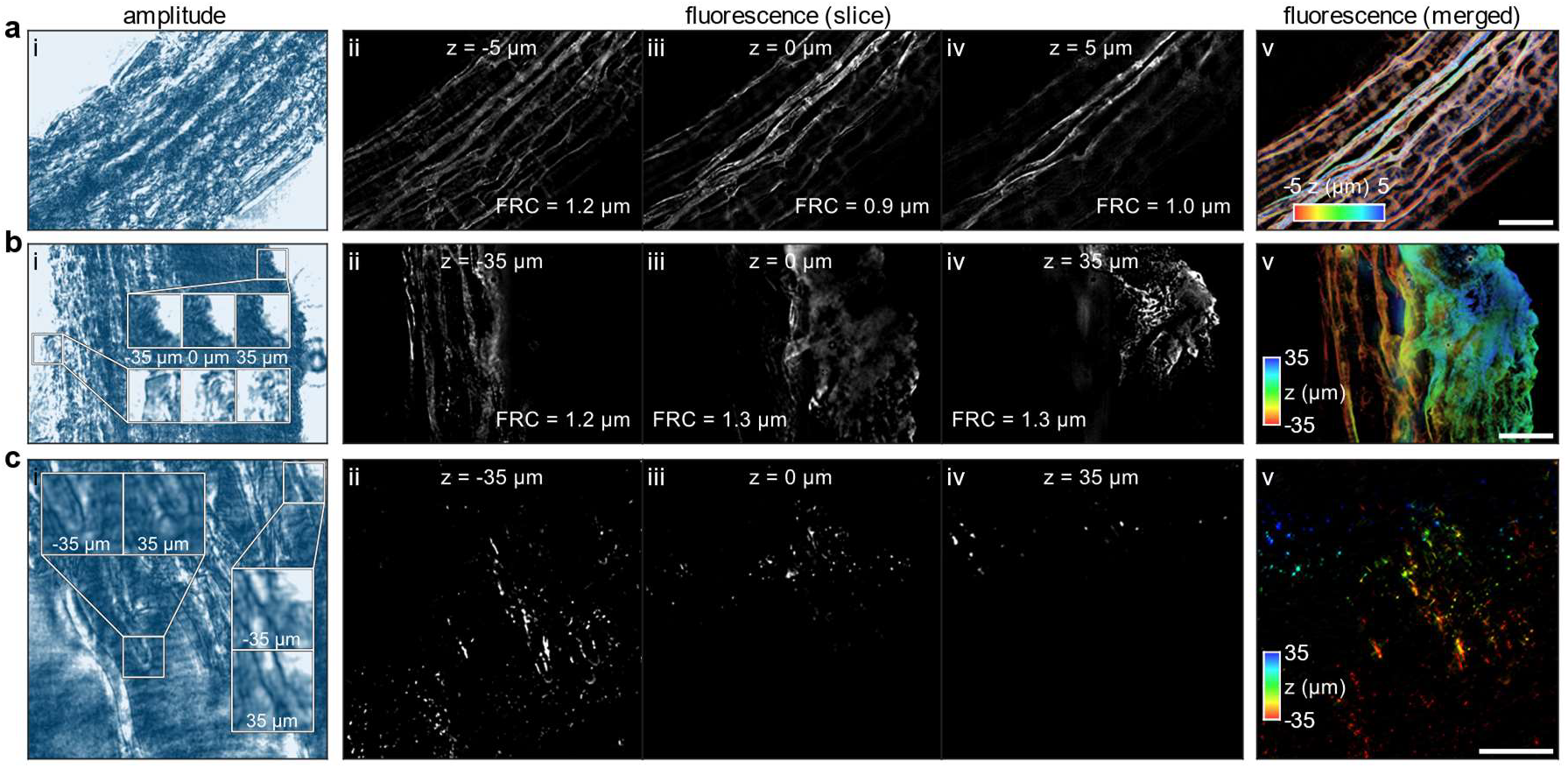
CFAST 3D imaging of maize roots. (a) A thin section and (b) a thick section on the root. (c) Imaging of fluorescent bacteria on thick section of roots. (i) Amplitude images, (ii-iv) reconstructed fluorescence images at various axial slices, and (v) 3D fluorescence images. Color bar, axial position. Scale bar, 50 microns.

We then test the ability of our system to capture bacteria on the surface of roots (i.e., the rhizoplane) with minimal acquisitions (Figure 2c and Supplementary Figure S5). Here, we use a grass called *Brachypodium distachyon* and a genetically tractable model rhizobacterium called *Pseudomonas synxantha* 2-79 (40) tagged with a constitutive reporter fluorophore (i.e., *P. synxantha* P_*PA10403*_*-mNeonGreen, P. syn-mNG*), which have been used to study root-microbe interactions previously (41). The thickness of the sample is obtained with amplitude and phase images at one axial point allowing us to deduce the number of captures required for visualizing the full volume of bacteria on the root surface. Again, considering a depth of field of 20 microns in 3D fluorescence imaging, we determine that four axial positions are required to obtain the 3D volume reconstruction of fluorescent bacteria on the root surface [Figure 2c(ii-v)].

### Bimodal imaging of bacteria early-stage colony biofilm formation

Roots typically contain surface-associated microbes called biofilms (42). As a proof of principle for studying biofilm development on roots, we demonstrate the utility of CFAST by imaging static bacterial aggregate development using a simple method to grow biofilms (43). Here, we track bacterial growth, competition, and gene expression during the early stages (i.e., within the first 24 hours of growth) of biofilm formation.

We first conduct timelapse imaging of *P. syn-mNG* on a MOPS based defined agar medium (Figure 3a,b). The bacterial surface area (i.e., radial growth in 2D) is captured by the fluorescence images [Figure 3a,b(i)]. The amplitude images [Figure 3a(iii)] show no morphological correlation with the fluorescence images [Figure 3a(i)], suggesting that the growth cannot be observed using a conventional brightfield microscope. In contrast, quantitative phase images [Figure 3a,b(ii)] closely mirror the fluorescence images [Figure 3a,b(i)]. The phase values within the biofilm area are easily distinguishable from the background (Figure 3c,d); the surface areas determined by the fluorescence and quantitative phase images exhibit a difference of less than 1.4% (Figure 3e). This experiment establishes that quantitative phase imaging can be used to study early biofilm development.

**Figure 3.**
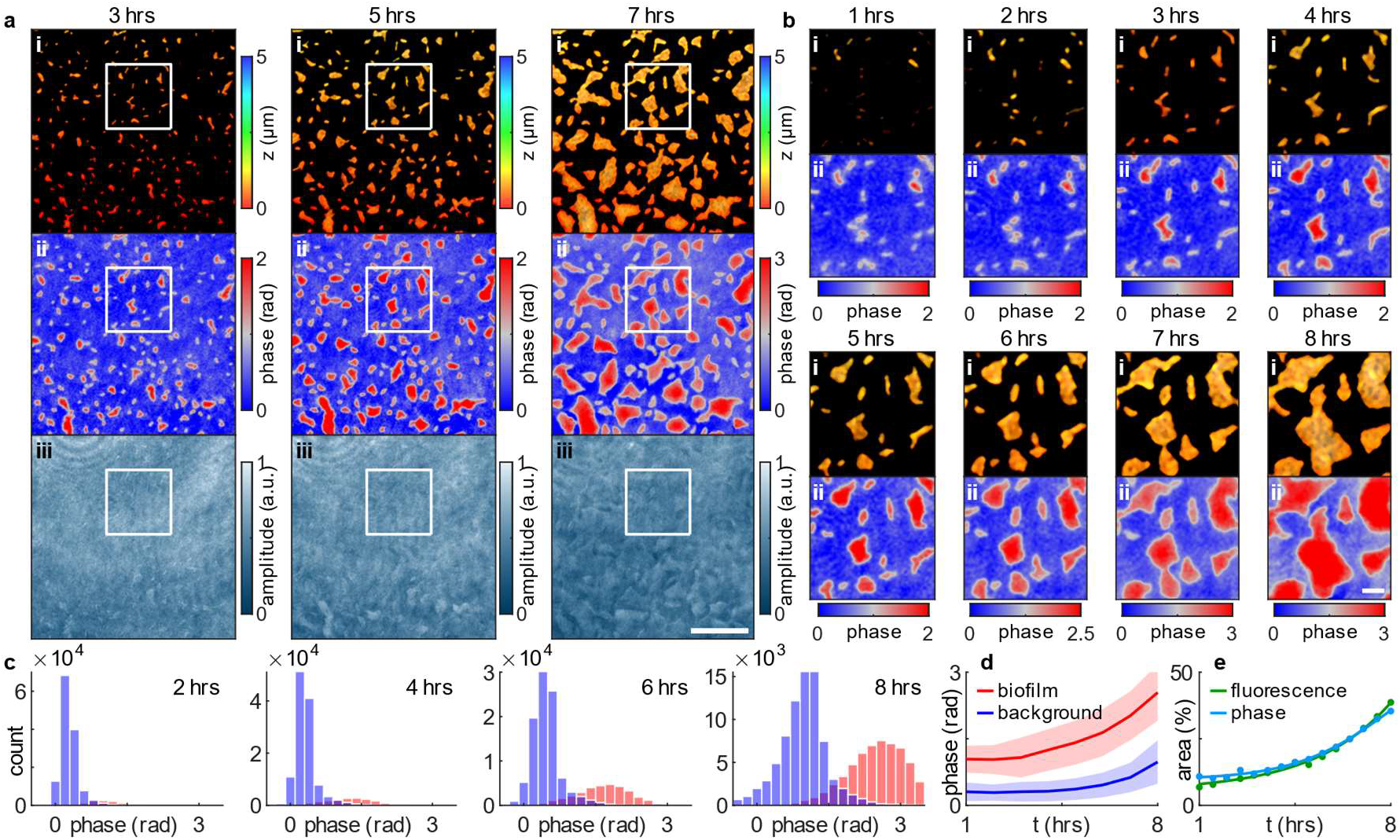
Time lapse CFAST imaging of bacterial biofilm formation. (a) Images of *P. synxantha* P_*PA10403*_*-mNeonGreen* (*P. syn-mNG*) growing on MOPS based defined agar medium. (b) The boxed region in (a). (i) Fluorescence images, (ii) quantitative phase images, (iii) amplitude images. Color bar, axial position in (i); phase in (ii); amplitude in (iii). Scale bar, 50 microns in (a); 10 microns in (b). (c) Distribution of phase values in the (red) bacterial biofilm and (blue) background areas. (d) Phase values of both areas over time. Solid lines represent the average; shaded areas represent the standard deviation. (e) Surface area of biofilm colony determined by the (light blue) quantitative phase images and (green) fluorescence images over time.

We next examine the ability of our method to investigate competition between *P. syn-mNG* and two wheat isolates that are not yet genetically tractable. Here, areas with fluorescence signal pinpoint the location of *P. syn-mNG*, while those with large phase values but no fluorescence identify the non-fluorescent strains (Supplementary Figure S6). We conduct a competition experiment with *P. syn-mNG* and *Paraburkholderia graminis* (*P. gram*), a recently collected wheat isolate (44). The initial concentration ratios tested are 1:5 and 1:20 (*P. syn-mNG* : *P. gram*) (Figure 4a). At the 1:5 ratio, *P. gram* is immediately outgrown by *P. syn-mNG* [Figure 4a(iv)]; it reaches stationary phase at ∼12 hours, occupying 5.9% of the total area. With the 1:20 ratio, *P. gram* reaches stationary phase at ∼13 hours, ultimately covering 16.0% of the total area [Figure 4a(iv)] yet is still outcompeted by *P. syn-mNG*. We then conduct a second competition experiment with *Pseudomonas orientalis* (*P. orien*) another wheat isolate. The initial concentration ratios tested are 1:5 and 1:1 (*P. syn-mNG* : *P. orien*) (Figure 4b). In this scenario, *P. orien* outcompetes *P. syn-mNG* at the 1:5 ratio and no competition is observed between the strain at a 1:1 ratio [Figure 4b(iv)].

**Figure 4.**
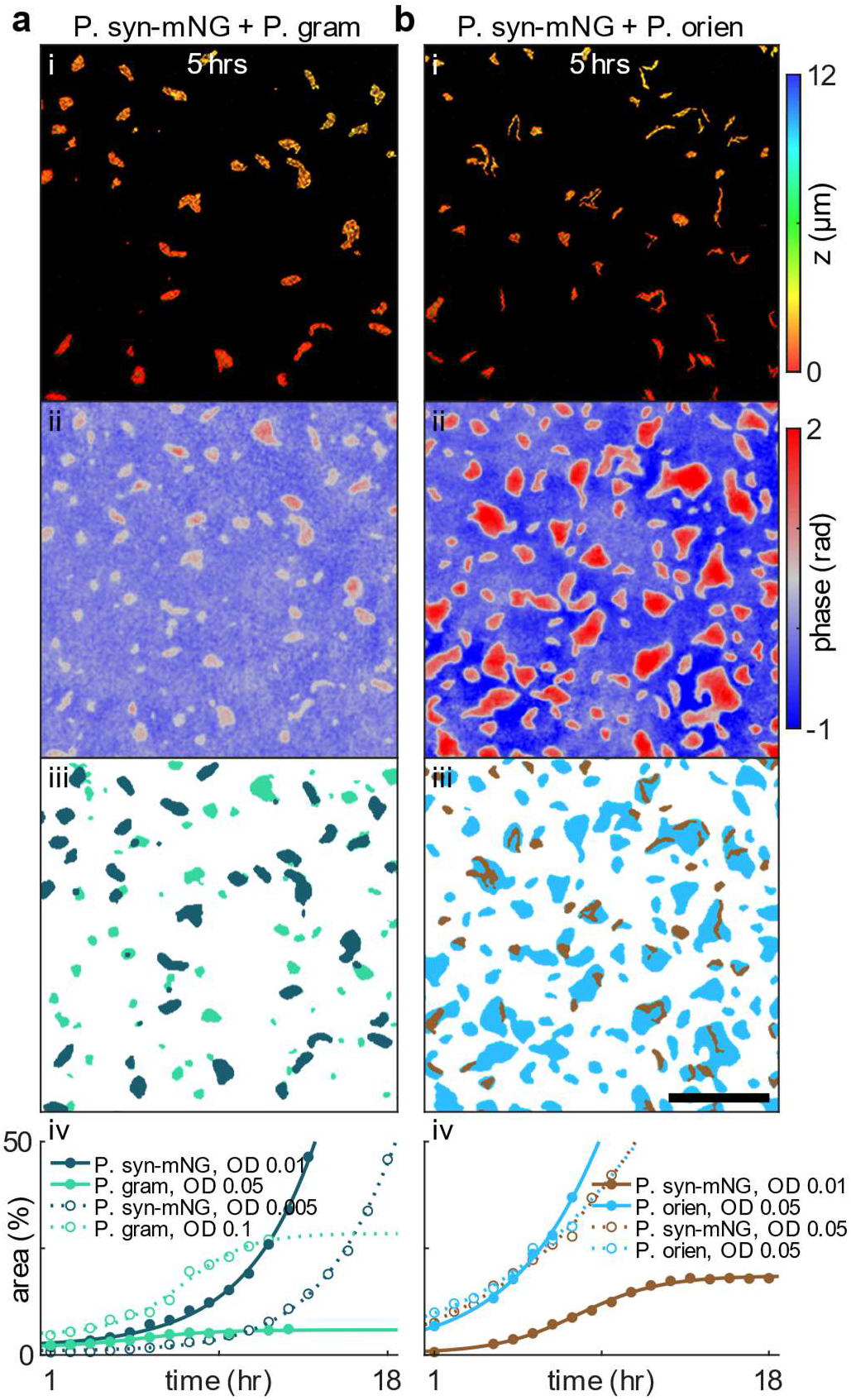
Differentiation of bacterial strains using CFAST. (a) Mixture of *P. syn-mNG* and *Paraburkholderia graminis* (*P. gram*) at 1:5 and 1:20 ratio. (b) Mixture of *P. syn-mNG* and *Pseudomonas orientalis* (*P. orien*) at 1:5 and 1:1 ratio. Representative images at the 5-hour timepoint showing that (i) fluorescence and (ii) quantitative phase images can be used to pinpoint (iii) different strains. Scale bar, 50 microns. Color bar, axial position in (i); phase in (ii). (iv) surface area percentage of different bacteria over time indicative of growth.

The 3D imaging capability of CFAST allows us to quantify spatiotemporal vertical growth dynamics during colony biofilm formation (Figure 5). Aggregate 3D growth kinetics are visualized over time along with the merging of aggregates into microcolonies (Figure 5a,b). We obtain the thickness probability at specific time points (Figure 5c) and a temporal progression of the thickness of all the aggregates and microcolonies within the field of view (Figure 5d). This result indicates that the thickness of the *P. syn-mNG* aggregates is greater when grown in the presence of *P. gram* rather than *P. orien*, potentially due to less competition and more nutrient availability. Moreover, the morphologies [Figure 5a,b(i)] of the *P. syn-mNG* aggregates with each competitor are strikingly different due to limitations and availability of space on the surface of the agar pad. Note that the 3D feature is not available for the quantitative phase images, therefore 3D images are not generated for non-fluorescent strains.

**Figure 5.**
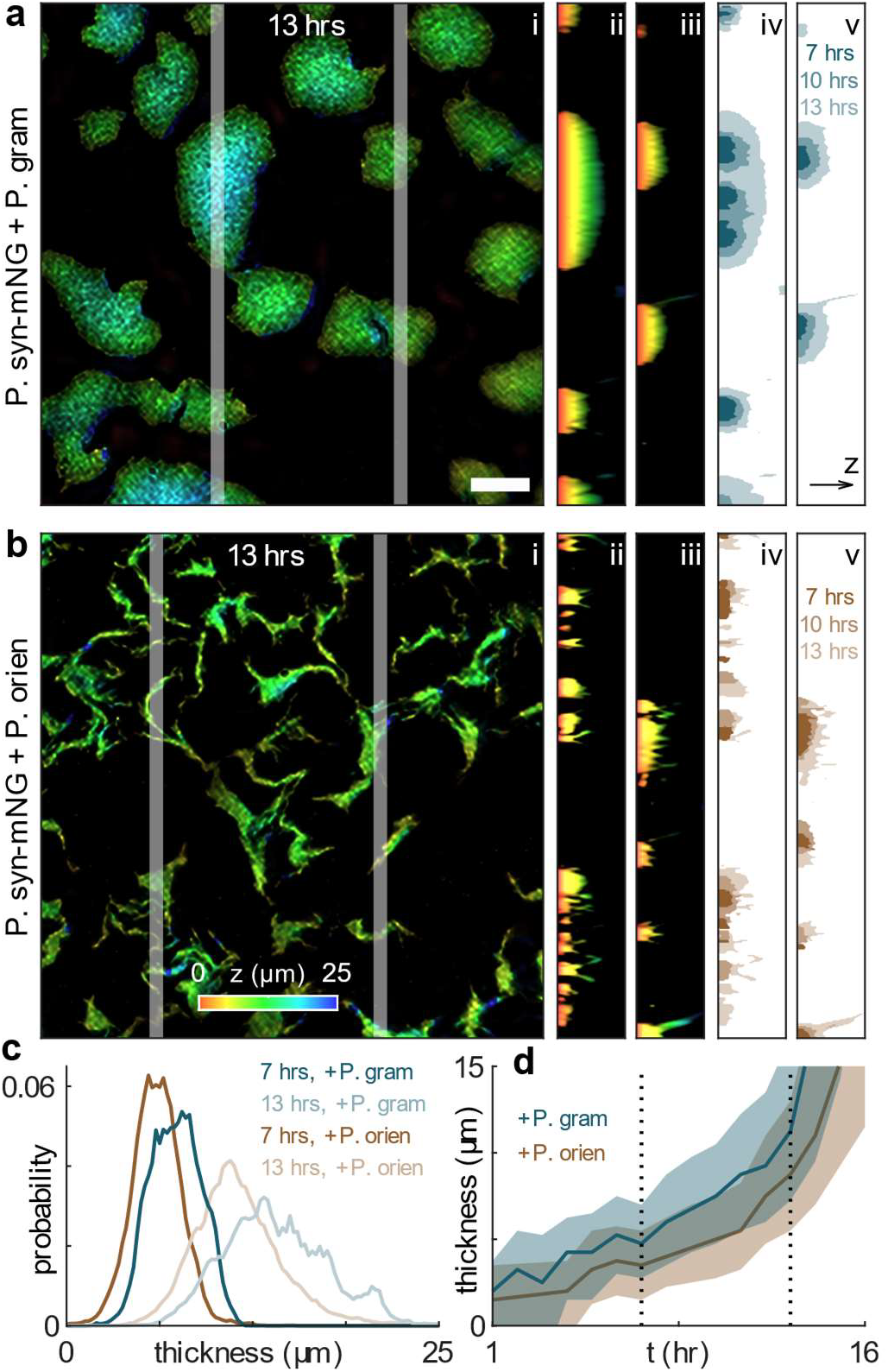
CFAST imaging of 3D growth dynamics during colony biofilm formation. (a) Mixture of *P. syn-mNG* with *P. gram*, (b) mixture of *P. syn-mNG* and *P. orien*. (i) The xy-view, (ii,iii) the yz-view, and (iv,v) the corresponding timelapse vertical growth (yz-view). Scale bar: 20 microns. Color bar: height in microns. (e) Thickness distribution of *P. syn-mNG* colony biofilm at 7 hours and 13 hours. (f) Thickness distribution over time. Solid lines represent the median; shaded areas represent the 33rd and 67th percentile.

Finally, we simultaneously image the growth with complex field and gene expression with fluorescence using CFAST. Here, we use transcriptional fusion reporters to monitor gene expression associated with promotors sensing phosphorus limitation (*P. synxantha* P_*phoA*_*-mNeonGreen, P. syn* P_*phoA*_*-mNG*) and controlling phenazine biosynthesis (*P. synxantha* P_*phzA*_*-mNeonGreen, P. syn* P_*phzA*_*-mNG*). In *Escherichia coli, phoA* is upregulated under phosphorus limitation (45), whereas the *phzA-G* operon in *P. synxantha* has been reported to be upregulated either by phosphorus limitation or at high-cell densities due to quorum sensing (46–49). We confirm these regulatory patterns with our constructs in planktonic conditions (Supplementary Figure S7). Surprisingly, these gene expression patterns do not always hold when bacteria are surface-attached. Figure 6a depicts the growth of *P. syn* P_*phoA*_*-mNG* and induction of P*phoA* under phosphorus limitation: induction occurs radially as the aggregate grows. The periphery of the aggregate has minimal induction, consistent with the idea that the periphery still has sufficient amounts of phosphate. Induction of P*phoA* only occurs under phosphorus limitation and correlates with batch conditions (Figure 6a,b,d, Supplementary Figure S7). However, induction of P*phzA* does not occur under phosphorus limited conditions but only after the surface area of the agar is uniformly covered by cells, indicating that induction under these conditions is due to quorum sensing only (Figure 6c,e). This result contrasts that of planktonic conditions, where induction occurs in response either to phosphorus limitations or quorum sensing (Supplementary Figure S7).

**Figure 6.**
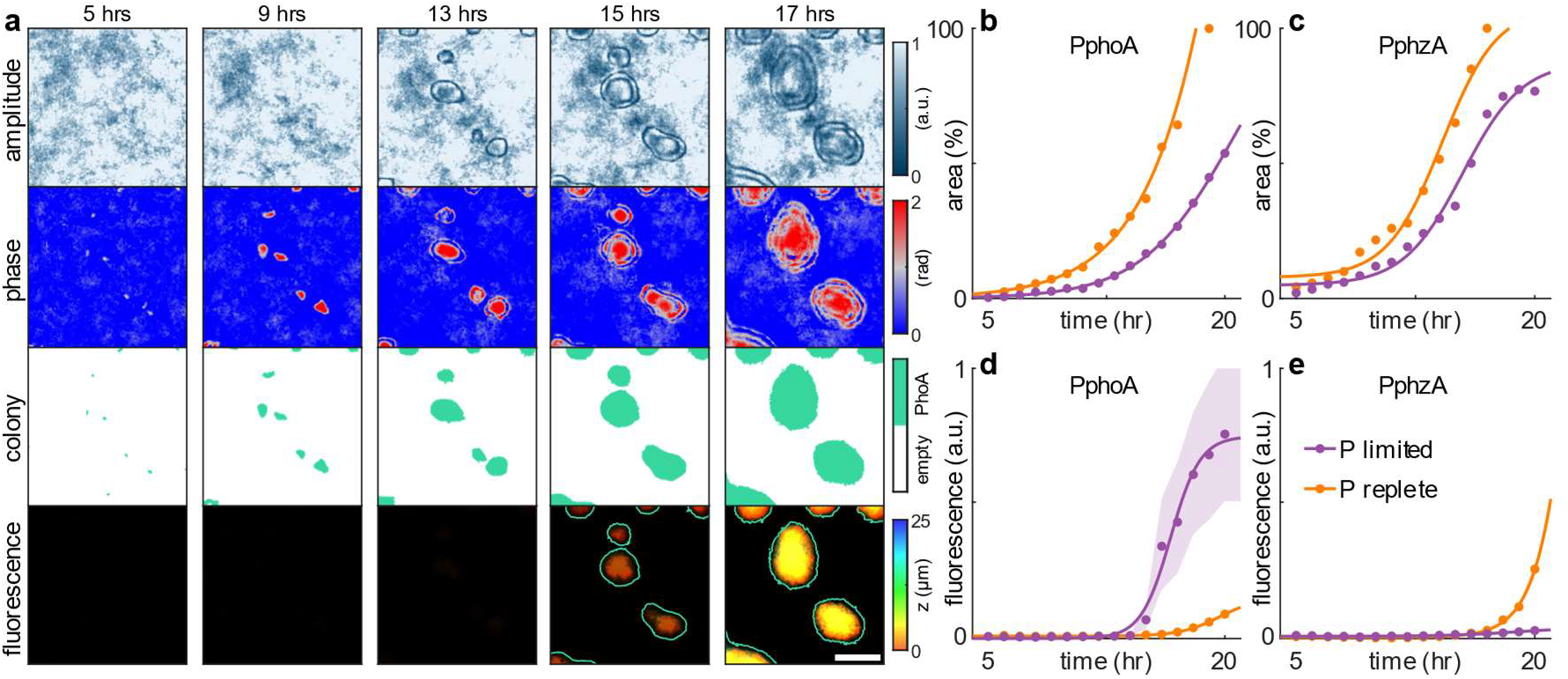
CFAST imaging of gene expression. (a) Imaging of *P. syn* P_*phoA*_*-mNG* under phosphorus (P) limitation. The growth is tracked by the complex field and gene expression by fluorescence. Scale bar: 50 microns. (b,c) Surface area percentage indicative of growth and (d,e) fluorescence level of the (b,d) *P. syn* P_*phoA*_*-mNG* and (c,e) *P. syn* P_*phzA*_*-mNG* colony biofilm over time. Shaded areas represent the 33rd and 67th percentile in (d,e). Purple: phosphorus limited condition; orange: phosphorus replete condition.

## Discussion

CFAST is a novel bimodal imaging technique that combines the strengths of both phase and fluorescence imaging. A key innovation of CFAST is the utilization of the spinning disk aperture, making it easy to integrate into existing commercial microscopes. Our results demonstrate that CFAST achieves lateral and axial resolutions of 0.6 and 1.9 microns, respectively. Employing CFAST, we capture 3D volumes of diverse roots morphologies with varying thicknesses, bacteria on roots, and bacterial biofilm gene expression and interspecies competitive dynamics in 3D.

We first showcase CFAST’s powerful depth of field when examining maize roots, enabling adaptive rather than exhaustive sample scanning to capture the 3D fluorescence distribution. This feature is critical when imaging samples of varying thickness as it minimizes potential photobleaching of fluorophores and phototoxicity to the sample, all while enhancing imaging speed. Moreover, we use the same technique to image bacteria on the surface of roots with minimal acquisitions. For perspective, the 70-micron thick sample we capture with four acquisitions at our axial resolution of 1.9 microns would require a minimum of 74 acquisitions with confocal microscopy when obtained with a 50% z-stacks overlap (optimal overlap). A 15% overlap would still require 44 image captures for similar resolution.

We then display CFAST’s phase sensitivity and fluorescence specificity by imaging colony biofilm formation as a proxy for biofilm growth on roots. The integration of quantitative phase and 3D fluorescence imaging enables us to conduct spatiotemporal monitoring of early stages of biofilm growth and morphogenesis, visualize competitive growth of bacteria, identify vertical growth dynamics, and monitor gene expression spatiotemporally under varying environmental conditions. To date, most biofilm imaging methods capture aspects of these capabilities for bacteria harboring fluorescent proteins (50–55) but rarely do they focus on the early stages of biofilm development, much less for mixed microbial communities including non-genetically-tractable bacteria. The 3D capacity offered by CFAST allows for auto-refocusing on the desired focal plane using only four frames (Supplementary Figure S8), minimizing the risk of excessive laser exposure. Moreover, the ability to capture the 3D volume of 20-µm depth with a single capture minimizes photobleaching and phototoxicity. This is crucial for long timescale imaging because we observe slower bacterial growth in laser-exposed regions (Supplementary Figure S9). Lastly, it is notable that induction of P*phzA* only occurs when aggregates and microcolonies have covered roughly 100% of the surface area with cells, and not in response to phosphorus limitation, unlike in planktonic culture. This unexpected result, consistent over several independent trials (Supplementary Figure S10), raises the possibility that certain regulatory dynamics may be affected by surface-sensing and motivates future studies using CFAST to reveal novel phenomena *in situ*, paving the way for follow-up mechanistic research.

In conclusion, CFAST is a valuable tool for imaging complex biological systems. Going forward, CFAST may be used to study how varying environmental conditions (be they from abiotic or biological sources) modulate microbial growth and gene expression on complex surfaces such as roots. Further, its efficiency in reconstructing full 3D volumes from a minimal number of images opens the possibility of real-time tracking the propagation of gene expression along the z-axis (i.e., through the thickness of a sample). Though the experiments reported here demonstrate CFAST’s utility for studying rhizosphere organisms in a laboratory setting, we envision alternations to CFAST’s design in the near future that would enable it to be buried underground, paving the way to *in situ* rhizosphere imaging.

## Materials and Methods

### CFAST forward imaging model

For complex field imaging of an object *s*(*x, y*), the optical field at the pupil plane modulated by the spinning disk is given by

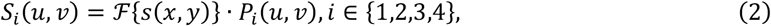

where *ℱ* represents the Fourier transform, (*x, y*) and (*u, v*) denotes the coordinates at sample and pupil planes, and *P*_*i*_(*u, v*) represents the binary amplitude mask defined by the *i*th spinning disk rotation angle. The camera then captures the intensity of the image plane, given by the square of the inverse Fourier transform of the modulated field *S*_*i*_(*u, v*),

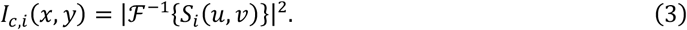

For 3D fluorescence imaging, the images ***I***_*f*_ *=* [*I*_*f*,1_, *I*_*f*,2_, *I*_*f*,3_, *I*_*f*,4_] are expressed as the convolution of an *N* -slice 3D object 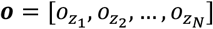and the PSF of the imaging system, 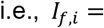 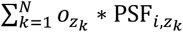, where 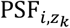 represents the PSF for the *i* th image at axial position *z*_*k*_ . For simplicity, we define an operator ***H*** that projects the object to the image space and rewrite the forward model as ***I***_*f*_ = ***Ho***.

### CFAST reconstruction algorithm

To reconstruct the object *s*_*i*_ from four intensity measurements *I*_*c,i*_, we first decompose the corresponding spectrum *S*_*i*_ into a scattered field 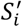 and an unscattered plane wave,

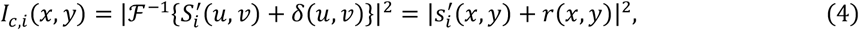

where 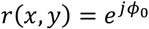 represents the plane wave and *ϕ*_0_ is a constant phase offset. Its Fourier transform is the Dirac delta function *δ*(*u, v*) corresponding to the normal incidence (NI) component in Figure 1b. To reconstruct the amplitude and phase of the optical field, we design an auxiliary function

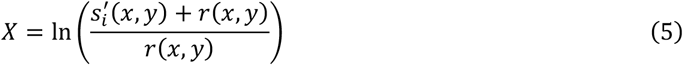

with a real part given by

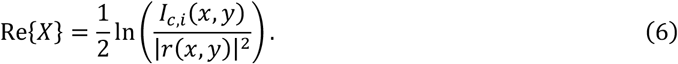

It has been shown that *X* is analytical in the upper half plane if the NI term is at the edge of the pupil and the unscattered light is stronger than the scattered light(29–31). Therefore, its imaginary part can be found using the Kramers-Kronig relation,

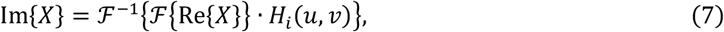

where Hilbert kernel *H*_*i*_(*u, v*) *=* −*j* sgn *u* for *i* ∈ {1,2} and *j* sgn *u* for *i* ∈ {3,4}. Modulated fields *S*_i_ is therefore given by

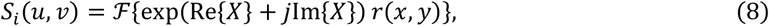

yielding the expression in Eq. (1).

To reconstruct the 3D object ***o*** from four detected fluorescence images ***I***_*f*_, we implement a modified Richardson-Lucy method to iteratively find the solution. The deconvolution algorithm can be written as

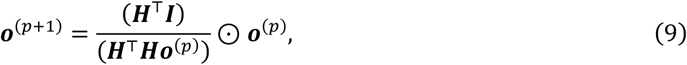

where ⊘ and ⊙ represent the element-wise division and product, respectively. Operator ***H***^T^ performs the backward projection 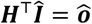where 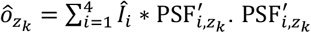 represents a 180°-rotated version of 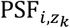.

### Maize and *Brachypodium distachyon* root preparation for bimodal imaging

#### Maize preparation

Maize seeds were obtained from Home Depot (cat. Number 78031-6). Seeds were first sterilized with 70% ethanol for 20 seconds, rinsed with autoclaved water 3x, placed in 50% bleach with 0.01% triton x for 3 minutes, and rinsed with autoclaved water 5x. Seeds were then pre-germinated on blot paper for 3 days. After germination, seedlings were transferred to 3 by 5-inch pots containing sterile sand. The maize plant was irrigated and watered with 0.5× Murashige and Skoog medium (0.5× MS medium, Sigma Aldrich, cat. Number M5524) adjusted to pH 5.8 with KOH. The plant was growth for roughly 3 months at room temperature near a laboratory window. Roots were then collected and cleared with 1M KOH overnight and rinsed with 0.1M HCl before imaging.

#### Brachypodium distachyon preparation

*Brachypodium distachyon* BD21 seeds were first sterilized with chlorine gas following previous protocols. Seeds were then plated 0.6% phytagel (Sigma Aldrich, cat. Number P8169) containing 0.5× MS and transferred to a growth chamber with a day/night cycle of 16hr/8hr at 25°C for 2 days. After 2 days of growth, the seedlings were dipped for 30 minutes into 0.5× MS buffer containing bacteria at an OD600 of 0.1. Finally, they were gently transferred to the surface of a MatTek plate (MatTek, cat. Number P35G-1.5-20-C) containing 0.6% phytagel with 0.5× MS medium and imaged shortly after.

### Bacterial strains and medium preparation

#### Strains

Six bacterial strains were used in this study. They include wild-type *Pseudomonas synxantha* 2-79 (gifted by Dmitri Mavrodi), *Pseudomonas synxantha* P_*PA10403*_*-mNeonGreen* (this study), *Pseudomonas synxantha* P_*phoA*_*-mNeonGreen* (this study), *Pseudomonas synxantha* P_*phzA*_*-mNeonGreen* (this study), *Pseudomonas synxantha* P_*tac*_*-mCherry* (gifted by Dmitri Mavrodi), *Paraburkholderia graminis* (wheat isolate), *Pseudomonas orientalis* (wheat isolate). Wheat isolates were collected from a non-irrigated wheat field at Washington State University’s Lind Dryland Research Station on August 9, 2019 (44).

#### Liquid medium

The strains were grown in a MOPS defined liquid medium (MOPS medium) containing: 4.1×10^-4^ M MgSO_4_·7H_2_O, 6.8×10^-4^ M CaCl_2_ ·2H_2_O, 1.6×10^-2^ M NH_4_Cl, 7×10^-3^ M KH_2_PO_4_, 5.55×10^-2^ M glucose as the carbon source, 1× MEM amino acids solution (Gibco, 11130051), and a modified version of Aquil trace metals (56) containing 1×10^-5^ M Fe and 1×10^-4^ M EDTA. The medium was buffered with 25 mM MOPS to pH 7 (pH adjusted with 1 M NaOH). For 96-well plate experiments with limited phosphate, the concentration of KH_2_PO_4_ was reduced to 50×10^-6^ M KH2PO4.

#### Solid agar medium

Agar plates consisted of MOPS medium with 1% noble agar. For experiments with limited phosphate, the concentration of KH_2_PO_4_ in the agar plates was reduced to 10×10^-6^ M KH_2_PO_4_. Two milliliters were pipetted into MatTek plates.

### Fluorescent reporter construction

The transcriptional fusion fluorescent reporters were constructed using the mini-Tn7 vector insertion protocol from Choi and Schwiezer (57). A broad host-range mini Tn7 vector was used for single-copy gene integration into the bacterial chromosomal at a neutral site located downstream of the highly conserved *glmS* site. We first cloned the desired inserts (i.e., promoters and mNeonGreen fluorophore) into the multiple cloning site of a pJM220 mini-Tn7 plasmid (Addgene, cat. Number 110559). Fragments were assembled using Gibson assembly (58) and electroporated into *E. coli* DH10B. Transformants were selected for on Luria-Bertani (LB) agar with gentamicin (20 μg mL^-1^). The presence of the correct construct was verified via PCR. Purified mini-Tn7 element DNA and pTNS1 helper plasmid (obtained from *E. coli* helper strains SM10/pTNS1) were then introduced into electrocompetent *P. syn* wild type strain via electroporation. Transformants were selected for on LB agar with gentamicin (20 μg mL^-1^). Correct constructs were confirmed by PCR, sanger DNA sequencing (Larage, Inc. Culver City, CA), and phenotype assays. See *Planktonic 96-well plate assays* for phenotype confirmation. A list of primers and plasmids used for each construct can be found in Supplementary Tables S1-S3.

### Planktonic 96-well plate assays

For all experiments, cells were initially inoculated from a -80°C glycerol stock on LB broth overnight (16-18 hrs) at 30°C shaking at 250 rpm. Cells were then washed 1× in MOPS medium that only contained 50 µM phosphate and plated onto a 96-well plate for an initial OD600 of 0.05. Each well contained 200 µL of media and 10 µL of inoculum. The plate reader (Tecan Spark 10M) was set to 30°C with orbital shaking (Amplitude 2.5 mm, Frequency 216 rpm). Optical density (OD600) was collected at 600 nm and mNeonGreen fluorescence was collected at an excitation wavelength of 485 nm and emission wavelength of 585nm. Data points were collected every 15 minutes for 24 hrs. Note that phosphorus replete wells contained 7 mM phosphate and phosphorus limited wells contained 50 µM phosphate.

### Bimodal biofilm imaging experiments

For all experiments, cells were initially inoculated from a -80°C glycerol stock on MOPS defined liquid medium overnight (16-18 hrs) at 30°C shaking at 250 rpm. Cells were then washed 1× in MOPS medium, diluted to an OD of 0.25 and grown for 4-5 hrs in MOPS medium. After this pre-growth, cells were washed 1× MOPS medium and diluted to an OD of 0.05, unless otherwise stated in the main text. Cells were then spot plated on the MOPS agar medium by pipetting 5µL of sample to the center of the MatTek plate. Plates were air dried for 1 hr with the lid off then transferred to the CFAST imaging platform containing a custom-made imaging chamber (Supplementary Figure S11) used to limit water evaporation from the agar. Time-lapse images were collected every hour for 24 hrs. Note that for the phosphorus limited experiments the 5 µL spotted cell sample contained 7mM phosphate while the agar medium it was spotted onto only contained 10 µM phosphate.

## Acknowledgments

We thank Dmitri Mavrodi for kindly providing wild-type *Pseudomonas synxantha* 2-79 and *Pseudomonas synxantha* 2-79 P_*tac*_*-mCherry*. We are grateful to the Resnick Sustainability Institute for enabling resources and financial support. R.E.A. was further supported by an NSF Postdoctoral Research Fellowship in Biology (Grant No. 2209379).

## Notes

### Competing Interest Statement

The authors have declared no competing interest.

